# Evaluating theories of neural information integration during visual search

**DOI:** 10.1101/2024.07.03.601936

**Authors:** Abe Leite, Hossein Adeli, Robert M. McPeek, Gregory J. Zelinsky

## Abstract

The brain routes and integrates information from many sources during behavior. A number of models explain this phenomenon within the framework of mixed selectivity theory, yet it is difficult to compare their predictions to understand how neurons and circuits integrate information. In this work, we apply time-series partial information decomposition [PID] to compare models of integration on a dataset of superior colliculus [SC] recordings collected during a multi-target visual search task. On this task, SC must integrate target guidance, bottom-up salience, and previous fixation signals to drive attention. We find evidence that SC neurons integrate these factors in diverse ways, including decision-variable selectivity to expected value, functional specialization to previous fixation, and code-switching (to incorporate new visual input).

## Introduction and Theoretical Significance

Living organisms constantly leverage many sources of information to generate adaptive behaviors. A grand challenge facing theoretical neuroscience is understanding how the neurons and circuits supporting such behaviors integrate this information. Mixed selectivity theory (Rigotti et al., 2013) argues that these neurons encode high-dimensional combinations of factors, rather than individual factors, enabling many possible linear decisions over a population of cells.

Within this framework, several models offer mechanisms by which neurons may flexibly encode multiple sources of information. **(1) Category-free mixed selectivity** (Raposo, Kaufman, & Churchland, 2014) predicts that neurons are sensitive to arbitrary and diverse combinations of factors that maximize downstream flexibility. **(2) Decision-variable selectivity** (Hirokawa, Vaughan, Masset, Ott, & Kepecs, 2019) predicts that neurons encode task-relevant intermediate variables, like the expected value of an action, rather than arbitrary mixtures. **(3) Neural code switching** (Shi et al., 2023; also subspace-dependent selectivity, eg Kaufman et al., 2022) predicts that neurons’ roles change to maximize efficiency when the context changes (via new sensory input or internal decisions). **(4) Functional specialization** (Yang, Joglekar, Song, Newsome, & Wang, 2019) is complementary to mixed selectivity and predicts the presence of neural circuits dedicated to computations used in many tasks, such that neurons in these circuits will encode only the variables relevant to their computations.

In this study, we take on the challenge of distinguishing between these models using neural recordings.

### Information Integration in SC during Search

We classify single neurons’ selectivity using a dataset of superior colliculus [SC] recordings from macaques performing a multi-target visual search task (Conroy, Nanjappa, & McPeek, 2023). This dataset is ideal for our question because eye movements (controlled by SC) are driven by multiple factors including top-down target guidance, bottom-up feature contrast, and history of previous fixation (Wolfe & Horowitz, 2017).

In each trial of this foraging-like task, the monkey sees a grid of disks where roughly 1/3 share a target color (e.g., green, with the rest being red distractors) and freely searches these disks until it fixates the one associated with reward. The monkey’s attention is thus guided by both top-down target-color guidance and bottom-up color contrast, as well as a need not to revisit previously searched disks. These are the three sources of information that we consider in our analysis. Spiking activity from a single SC cell per session was recorded and aligned on fixation and saccade onset, and grids were generated such that this cell’s receptive field (RF) would fall on a disk adjacent to the currently-fixated disk. This allowed for a precise measure of how the recorded cell’s firing rate reflects the properties of the stimulus in its RF.

## Methods

We applied partial information decomposition [PID] to timeseries smoothed firing rate data from Conroy et al. (2023) to determine which theoretical model best fits each neuron, based on its encoding of previous fixation, target guidance, and contrast salience. The first two factors were labeled in the dataset, and we defined the last by whether most of a disk’s neighbors are a different color (roughly the median contrast).

In the interest of brevity, we refer readers to Williams and Beer (2010) for the details of the PID analysis, and clarify here only our application of PID. We performed separate PID analyses at each timestep and binned data into quartiles, such that a neuron active in more than 25% of trials would have a capacity of 2 bits. We used *I*_*min*_ as our measure of redundant information, and filtered the output with a .05 Monte Carlo significance threshold (understanding that this is not a formal significance test due to the lack of multiple test correction). We ran 1000 Monte Carlo runs, each of which shuffled firing rates over trials (keeping combinations of factors intact).

PID outputs 23 terms representing the extent to which a cell encodes each redundant, unique, and synergistic combination of the task variables. Redundant information is available whenever any source variable is available; synergistic information is available only when all are available; and unique information is available when and only when a particular variable is available. To help visualize these 23 terms, we represent each source variable as a primary color, and their combinations as darkened or lightened mixtures of the colors.

We classified cells to models using a hand-tuned algorithm based on visual inspection of the PID plots. This algorithm operates on each cell as follows, terminating when the cell is labeled: (1) mark all unique/pairwise synergistic terms whose areas (in the plot) exceed 0.25 bit ·ms. (2) if any pair of marked terms has an overlap of areas (in the plot) below 60%, label the cell as **code switching**. (3) if exactly one unique term (and no synergistic term) is marked, label the cell as **specialized**. (4) if the synergy between previous fixation and target guidance (representing expected value) is marked, label the cell as **decision-variable**. (5) if any term(s) are marked, label the cell as **category-free**. (6) fail to label the cell.

## Results

We performed time-series PID on 113 neurons across 2 monkeys, yielding rich signatures of the cells’ roles during the task, and we used these as described above to classify neurons. We found **4 category-free** cells, **13 decision variable** cells, **13 code-switching** cells, **25 specialized** cells, and 58 cells that could not be categorized. Of the 25 cells specialized on this task, 16 were specialized to previous fixation, 7 to feature guidance, and 2 to contrast salience. We present the strongest exemplars of each group in Figure 1.

**Figure 1:**
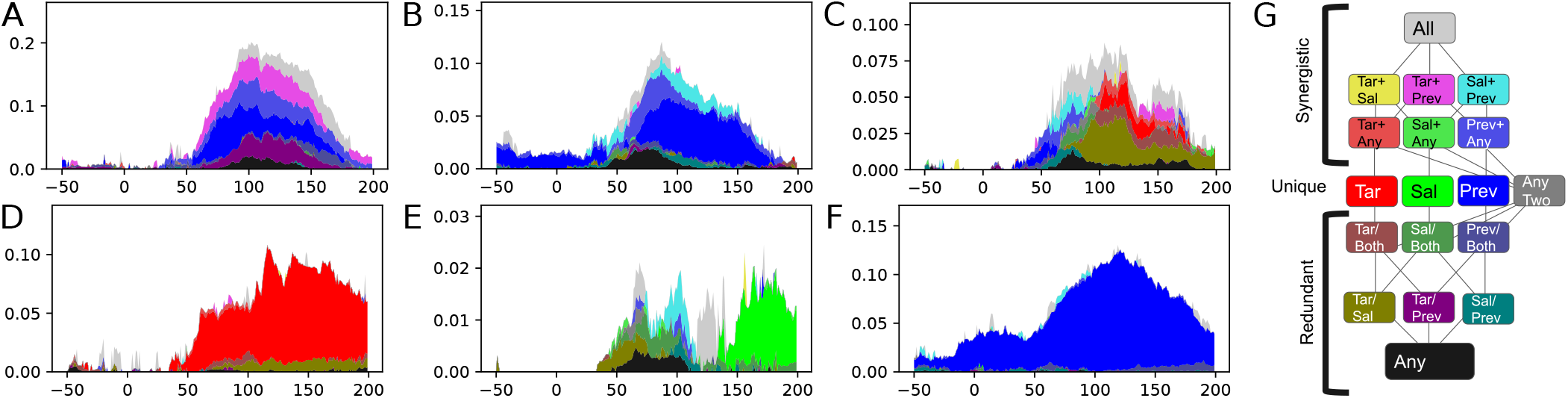
PID plots for neurons most consistent with each model: (A) decision-variable, (B) category-free, (C) code-switching, and specialization to (D) target guidance, (E) contrast salience, and (F) previous fixation. Information terms (in bits) are vertically stacked on the *y*-axis using color key (G); time from fixation onset is shown on the *x*-axis. (Must be viewed in color.)

The code-switching cell shown in Fig 1C illustrates the explanatory power of PID. This neuron initially encodes previous fixation (25-75ms); it adds contrast salience at 50ms, but this becomes redundant with target guidance by 100ms, with guidance and previous fixation showing a late synergy emerging at 125-175ms. This cell’s PID signature is compared in Fig 2 to single-variable and nested time-series GLM analyses and mean firing rate by condition. Consistent with PID, GLM conveys the redundance of contrast salience by its greater prominence in Fig 2A than 2B, and it conveys the late synergy of previous fixation by its prominence in Fig 2B but not 2A. The mean firing-rate plot is useful to elucidate effect directions, but is difficult to interpret on its own.

**Figure 2:**
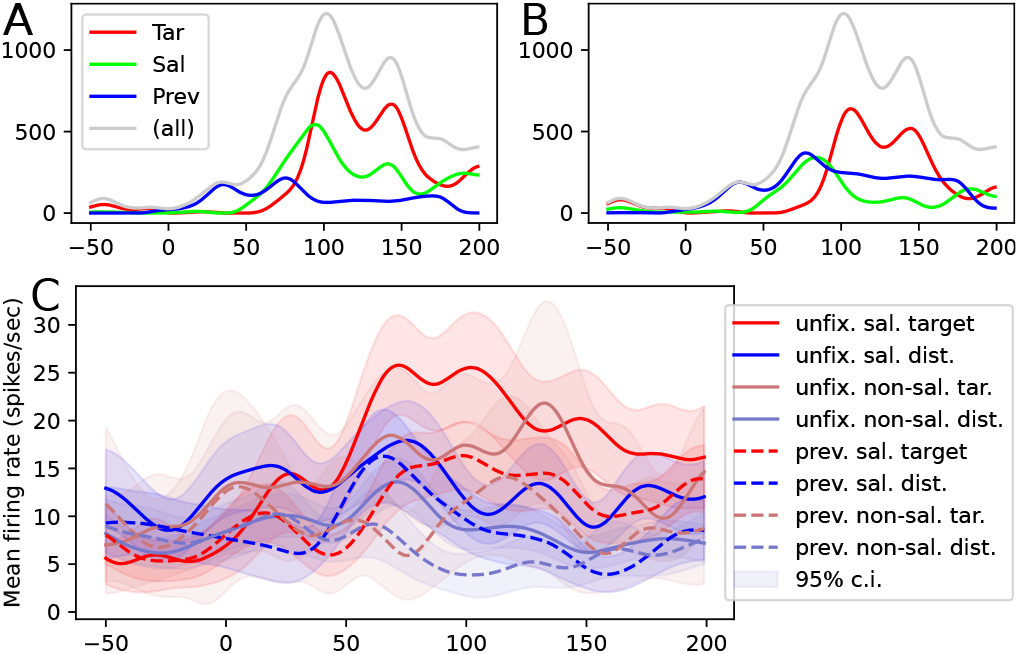
(A) single-variable & (B) nested time-series Poisson GLM plots of code-switching cell [Fig 1C], with log-likelihood delta on *y*-axis. (C) mean firing-rate plot over the 8 conditions.

## Discussion

### PID as a tool for visualizing neural representation

Compared to mean firing rate analyses and GLMs, time-series PID seems the best tool for this job. One needs a time-series analysis to evaluate hypotheses about neural code switching, and PID conceptually unifies the two common flavors of GLM: single-variable GLM measures ‘sufficient variance’ (redundant + unique information), and nested GLM measures ‘necessary variance’ (unique + synergistic information). Unlike GLMs, PID captures unique information and separates 2-way and 3way interactions. Additional advantages are that PID reports interpretable quantities relative to a cell’s information capacity (vs model likelihoods), and that it is robust to bimodal encoding of variables. Disadvantages of PID include its reliance on discretization, which reduces sensitivity, and an explosion of terms that limits its use to three factors.

### The SC is more than just a priority map

From the diverse neural signatures we report, it is clear that the SC is the site of meaningful integration across multiple sources of information; it does not merely encode a single priority map computed elsewhere. A neuron encoding priority would be classified as decision-variable in our analysis, but only 13 of the 55 categorized cells played this role. SC’s local computations include neural reuse, with 13 neurons changing their codes over fixation. Lastly, 16 neurons were specialized to previous fixation, suggesting future study of whether the SC includes a circuit dedicated to this computation. However, SC is not entirely flexible like PFC: only 4 cells encoded arbitrary combinations of task variables as predicted by category-free selectivity. Overall, our findings make it clear that the SC performs many computations relevant to attention and target selection locally, using a variety of methods to support the flexible visual behaviors it generates.

## Acknowledgments

This work was supported in part by NSF-GRFP award 2234683 to A.L. and by NIH-NEI award R01EY030669.

